# The Selenoproteome as a dynamic response mechanism to oxidative stress in methanogenic communities

**DOI:** 10.1101/2023.08.14.549203

**Authors:** Hugo Kleikamp, Paola A. Palacios, George Papaharalabos, Anders Bentien, Michael V. W. Kofoed, Jeppe L. Nielsen

**Author notes:** Corresponding author: Hugo Kleikamp.

## Abstract

Methanogenesis is a critical process in the carbon cycle that is applied industrially in anaerobic digestion and biogas production. While naturally occurring in diverse environments, methanogenesis requires anaerobic and reduced conditions though varying degrees of oxygen tolerance have been described. Micro-aeration is touted as the next step to increase methane production and improve hydrolysis in digestion processes; therefore, a deeper understanding of the methanogenic response to oxygen stress is needed. In order to explore the drivers of oxygen tolerance in methanogenesis, two parallel enrichments were performed in an environment without reducing agents and in a redox-controlled environment by adding redox mediator AQDS. The cellular response to oxidative conditions is mapped using proteomic analysis. The resulting community showed remarkable tolerance to high-redox environments and was unperturbed in its methane production. Next to expressing pathways to mitigate reactive oxygen species, the higher redox potential environment showed an increased presence of selenocysteine and selenium-associated pathways. By including sulfur-to-selenium mass shifts in a proteomic database search, we provide the first evidence of dynamic and large-scale incorporation of selenocysteine as a response to oxidative stress by methanogenic organisms and the presence of a dynamic selenoproteome.

## Introduction

Methanogenesis is a critical process in the carbon cycle in soils, anaerobic digestion processes and the gut microbiome, where it produces CH_4_ and prevents the accumulation of H_2_. It is responsible for harmful greenhouse gas emissions and beneficial biogas production processes. Methanogens are present in highly diverse habitats, including extreme environments (Jones 1983, Thauer 2008, Sorokin 2015), but generally require reduced environments and anaerobic conditions. Methanogens display varying degrees of oxygen sensitivity (Kiener 1983, Fetzer 1993, Kato 1993) and are typically cultured in the presence of reducing agents such as cysteine, thioglycolate or FeS, that can scavenge oxygen (Plugge 2005).

Large-scale genomic analysis has categorized methanogenic adaptation mechanisms to oxidative stress (Lyu 2018). Methanogens were shown to possess enzymes that could remove reactive oxygen species (ROS). These include catalases, peroxides, superoxide dismutases, nitric oxide reductases, which can directly eliminate ROS, as well as thioredoxin and rubredoxin, which provides redox buffering and the ferritin system for iron storage (Lyu 2018). Oxygen sensitivity has been linked to the Class I (*Methanobacteriales, Methanococcales, Methanopyrales*) and Class II (*Methanocellales, Methanomicrobiales, Methanosarcinales*) groups (Brochier-Armanet 2011). While both groups possessed ROS-mitigation pathways, Class II is often abundant in alternating oxic environments and contains more genes dedicated to ROS elimination and damage repair (Lyu 2018).

While ROS are able to damage DNA and metalloenzymes (Imlay 2013), methanogens are especially prone to oxidative damage due to their reliance on cysteine residues, which is a crucial component in the active sites in methanogenic enzymes (Klipcan 2008) and is found in both iron-sulfur clusters and Ni-Fe hydrogenases (Hamann 2007, Vitt 2014). Since oxidative stress can irreversibly oxidize cysteine to cysteic acid (Rabilloud 2002), another strategy to confer oxygen tolerance is using selenocysteine (Evans 2021), which, as opposed to cysteine, can readily reverse its oxidative damage (Maroney 2018). Selenocysteine, often referred to as the 21st amino acid, substitutes a thiol for a selenol group. Although low abundant, selenoproteins are found in all domains of life and are incorporated through an alternative translation of UGAs stop codon, a selenocysteine specific tRNA, and a neighboring selenocysteine insertion sequence (SECIS) element in the mRNA secondary structure (Barry 1991). In eukaryotes, selenoproteins play a role in oxidative stress as thioredoxin reductase and glutathione peroxidase (Rayman 2000). It has also been detected in relation to methanogenesis, where it is found in various hydrogenases (Rother 2010, 2018). Compared to its cysteine counterpart, selenocysteine confers different properties, such as lower pKa and redox potential, while also increasing nucleophilicity and enzymatic turnover (Axley 1991, Arnér 2010). Despite its benefits in methanogenesis, selenium utilization has only been identified in a limited number of archaeal genera (*Methanococcales, Methanopyrales*, and *Lokiarchaeota*) (Zhang 2022). However, the recoding of *Asgardarchaeota* stop-codons showed that they could be much more widespread in this domain (Liu 2018, Sun 2021).

Deciphering the response of methanogens to oxygen and the role of selenocysteine will aid the understanding of biogeochemical cycles and the design of more robust and efficient digestion and biogas production processes (Muller, 2022, Shanfei, 2022). Therefore, in this study, we present a proteomic analysis of methanogenic enrichment cultures grown in the presence and absence of reducing agent AQDS (9,10-anthraquinone-2,7-disulfonate disodium), While the oxidation-reduction potential (ORP) was monitored with probes, proteomic analysis was used to examine oxidative stress and selenium-associated changes in the proteome.

## Materials and Methods

### Sampling and Culture Conditions

The sludge was collected from a mesophilic (39 °C) anaerobic digester fed with organic residues and manure as substrates from the Bånlev Biogas plant (DK). The sludge was used as inoculum in a batch process supplemented daily with a H_2_/CO_2_ gas mixture (80:20, v/v) at an initial pressure of 1.8 bar. After one week of enrichment at 37 °C, the sludge was transferred to mineral phosphate medium containing (per liter): 5.0 g of K_2_HPO_4_, 2.5 g of KH_2_PO_4_, 1.0 g of NH_4_Cl, 0.05 g of KCl, 1.0 g of NaCl, 0.05 g of CaCl_2_·2H_2_O, 0.16 g of MgCl_2_·6H_2_O, 10 mL of 141 modified Wolin’s mineral solution (DSMZ), and 10 mL of 141 Wolin’s vitamin solution (DSMZ). Five consecutive transfers were performed afterwards (every two weeks), using 10 % inoculum, the mineral phosphate medium, and H_2_/CO_2_ (80:20, v/v) as the only energy and carbon source.

### Experimental set-up

Thirty milliliters of enriched culture from the 5^th^ transfer were inoculated into thirty milliliters of fresh mineral phosphate medium, for a working volume of 60 mL, in 155 mL reactors. The experimental set-up included four parallel reactors, divided into two different conditions: Two reactors contained 25 mM of oxidized 2,7-AQDS (95%), and two other reactors, in which 2,7-AQDS was omitted. Each reactor was equipped with an oxidation-reduction potential (ORP) probe (Atlas Scientific LLC, LongIsland City, USA) and a pH probe (Atlas Scientific LLC, LongIsland City, USA) for the continuous monitoring of redox potential and pH. The reactors were re-flushed daily with H_2_/CO_2_ gas mix (80:20, v/v) and pressurized to 1.8 bar. Before the daily re-flushing, the pressure of each reactor was measured.

### Metaproteomics

Samples of duplicate cultures without reducing agent (H-ORP) and the reduced environment with redox mediator 2,7-AQDS (L-ORP) were taken on days 21 and 22 of the enrichment for proteomics analysis. Cell lysis was performed with a Tissue Homogenizer (Covaris). Subsequent tryptic digestion, reduction and alkylation steps and peptide purification were performed using the Its kit (PreOmics). ESI LC-MS was performed on a QE Orbitrap system (Thermo Scientific) with a 120-minute gradient (García-Moreno 2020).

### Metaproteomics data analysis

Database searching was performed against a sample-guided GTDB r207 database (Parks 2022) with the CHEW pipeline (github.com/hbckleikamp/CHEW), using oxidation, phosphorylation, acetylation and pyroglutamate formation as variable modifications. An additional variable modification corresponding to the mass shift between sulfur and selenium (S-Se) was included for cysteine and methionine. Matched peptides were filtered using a reversed decoy database and a 0.05 false discovery rate. Pathways related to ROS mitigation were composed of either EC numbers, KEGG orthologies or InterPRO groups (STable 1). Functional annotation was performed with GhostKOALA to obtain KEGG orthologies (Kanehisa 2000, 2016) and with homology search against UniprotKB with DIAMOND (Buchfink 2015) to obtain EC numbers and InterPRO groups. To identify selenoproteins, which are absent in GTDB, an additional database search was performed against UniprotKB reference proteomes corresponding to species identified in the initial search against GTDB.

## Results

### Enrichment under oxidative conditions

Two samples of enriched full-scale mesophilic sludge were incubated in the presence and absence of 25 mM of reducing agent AQDS. Due to under-pressure from H_2_ and CO_2_ consumption, irregular and short exposure to small amounts oxygen occurred during the entire incubation period (Figure 1A) until the headspace gas was replaced. This resulted in culture conditions with fluctuating and higher ORP (H-ORP), while the enrichment grown in the presence of AQDS showed a much more stable ORP, and lower redox potential (L-ORP) (Figure 1A). In the first days, the ORP decreased from -200 to -400 mV as oxidized AQDS was reduced to AQDSH_2_ (Figure SI 2A), after which the L-ORP culture was able to maintain a stable median ORP of -342 and -378 mV, which is close to a previously described optimum of -351 mV (Vongvichiankul 2017). Proteomic sampling occurred on days 20 (a) and 21 (b), at which both cultures were sufficiently adapted and showed similar methane yields (Figure SI 2B).

**Figure 1.**
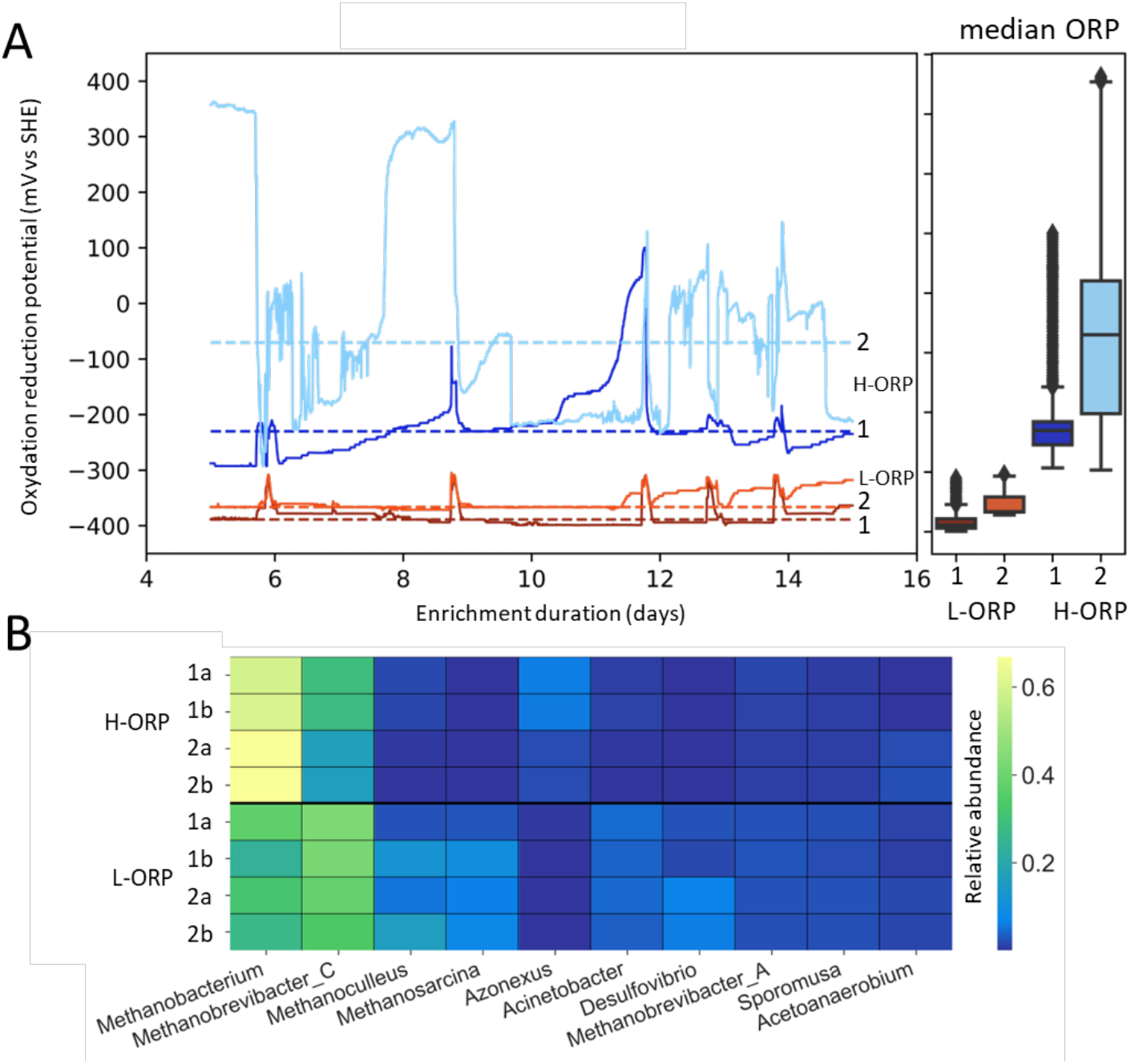
A: oxidation-reduction potential (ORP) graphs of replicate (1,2) methanogenic enrichments grown with 25 mM of redox mediator AQDS (L-ORP, orange) and enrichment without reducing agents (H-ORP, blue) during an enrichment period. Samples for proteomic analysis were taken on days 20 (a) and 21 (b). Median ORP values for each culture are shown with dotted lines and boxplots. B: taxonomic abundance normalized to the top 10 most abundant genera.

Although one H-ORP reactor reached redox potentials above 200mV, well above the preferred redox environment for methanogenesis, methane yield was not affected. This shows that this enrichment possessed significant tolerance towards temporary oxidative stress events. Metaproteomic analysis of both communities revealed that the H-ORP condition was highly enriched in *Methanobacterium* (68 %) and *Methanobrevibacter_C* (16 %), both Class I methanogens, classified as oxygen sensitive, and had lower diversity in methanogens compared to the enrichment in L-ORP, which, next to *Methanobrevibacter_C* (39 %) *Methanobacterium* (30 %), had more *Methanoculleus* (9 %) and *Methanosarcina* (6,5 %), which are part of more oxygen tolerant Class II methanogens (Figure 1C). Both samples retained the ability to produce methane and highly similar functional profiles, dominated by methane metabolism (Figure SI 3).

### Expression of ROS mitigation pathways

Functional annotation was performed to investigate the physiological response to oxidative stress using a combination of EC numbers, KEGG pathways and InterPro families (Table SI 1). Various enzymes and protein families associated with ROS mitigation were found to be expressed, such as catalases, peroxidases and superoxide dismutases, as well as thioredoxin and rubredoxin-associated proteins (Figure 2A). Clustering of expression profiles across the two enrichments with and without AQDS showed that most ROS-mitigating protein functions clustered together and were more expressed in the H-ORP samples (Figure 2A). In addition, they grouped with selenium-associated proteins and cysteine and methionine metabolism. Next to ROS mitigating proteins, pathways involved in repair of ROS damage were analyzed, including base excision repair and biogenesis of iron-sulfur proteins, but these showed different expression patterns. Detected selenium-associated proteins included a putative selenium binding protein YdfZ, cysteine desulfurase SufS and SepSecS/SepCysS tRNA synthase, which are linked to selenocysteine, as well as selenium transferase SelA and elongation factor SelB, which are key enzymes involved in selenocysteine incorporation (Figure 2B).

**Figure 2.**
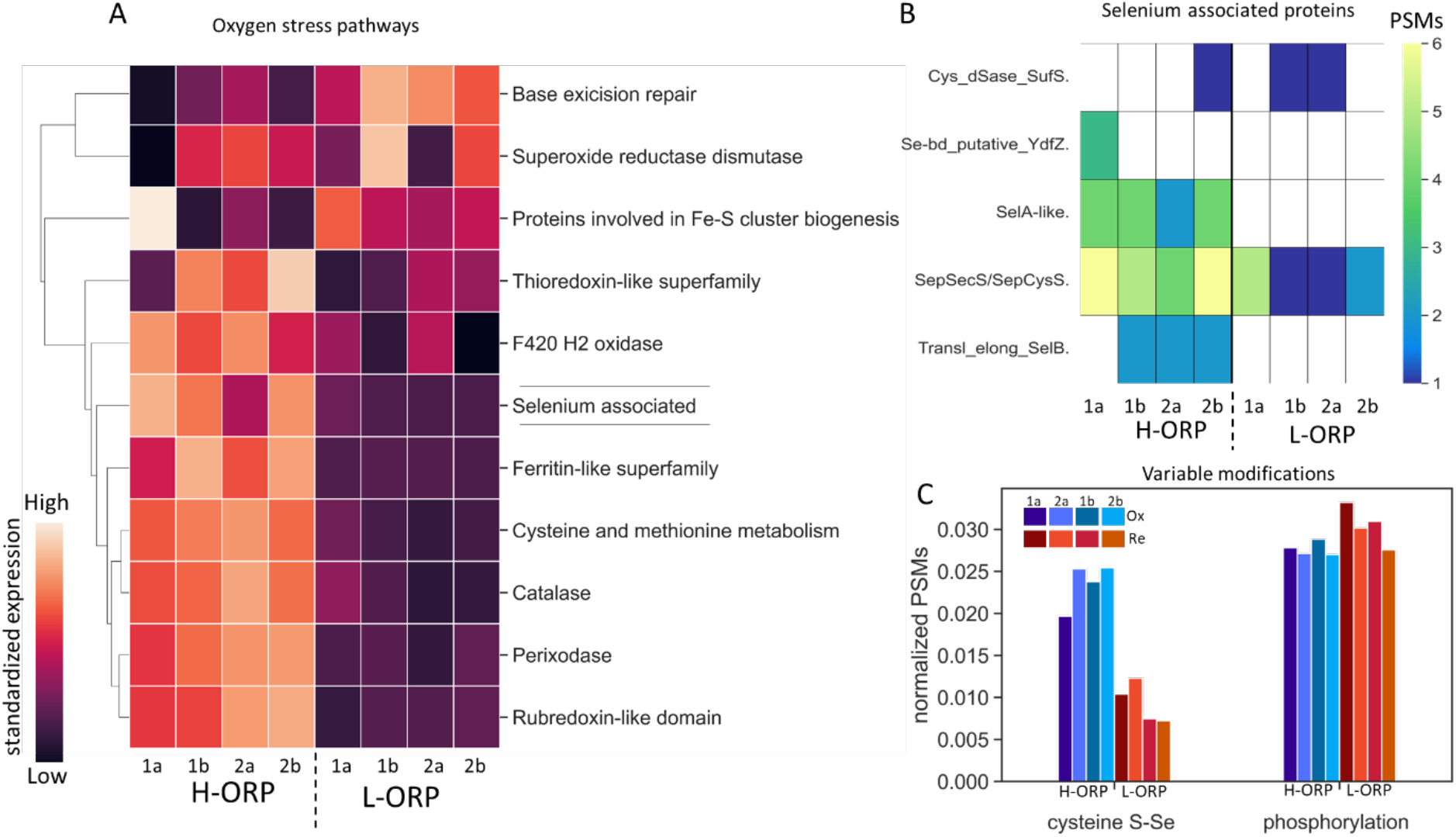
A: cluster map of expression of enzymes, pathways, and protein families associated with oxygen stress. Normalized peptide spectrum matches were standardized across samples from two enrichments without reducing agents (H-ORP) and grown with 25mM of redox mediator AQDS (L-ORP), and clustered based on Euclidean distance to relative shifts in expression. Higher expression is indicated by bright coloring versus dark in the case of lower expression. B: number of peptide spectrum matches (PSMs) that match selenocysteine-associated proteins, C: Normalized abundance of peptide spectrum matches (PSMs) of the S-Se mass shift compared to the abundance of phosphorylated peptides.

### Presence of selenocysteine

While selenium-associated proteins were identified in each sample, no selenoproteins were identified for archaea when using UniprotKB reference proteomes as an alternative protein database, which only identified bacterial selenoproteins in *Acetoanarobium, Desulfovibrio* and *Rubrivivax* (Figure SI 6). To analyze the presence of putative selenium incorporation, the S-Se mass shift, which corresponded to the mass difference between sulfur and selenium, was included as a variable modification in a proteomic database search to identify both selenocysteine and selenomethionine. The frequency of this S-Se shift was compared to other common protein modifications (Figure SI 4). The S-Se mass shift was observed consistently throughout both enrichments but was two and a half times more prevalent in the H-ORP enrichment and displayed similar identification rates to phosphorylation (Figure 2C). The S-Se mass shift was detected exclusively on cysteine residues, not methionine.

To analyze the taxonomic distribution of S-Se substitutions, the ten most abundant genera were clustered according to their relative abundance of all peptides and their number of selenocysteine-containing peptides. The abundance values of the genera were standardized across samples to enable clustering of abundance patterns and reveal underlying trends between the two samples with three distinct groups (Figure 3A). *Methanobacterium* displayed higher abundance of peptides and selenocysteine in the H-ORP enrichment, *Methanobrevibacter* A & C increased peptides and selenocysteine at low ORP, while Group III consisted of several genera with increased abundance at low ORP but less change in selenocysteine incorporation.

**Figure 3.**
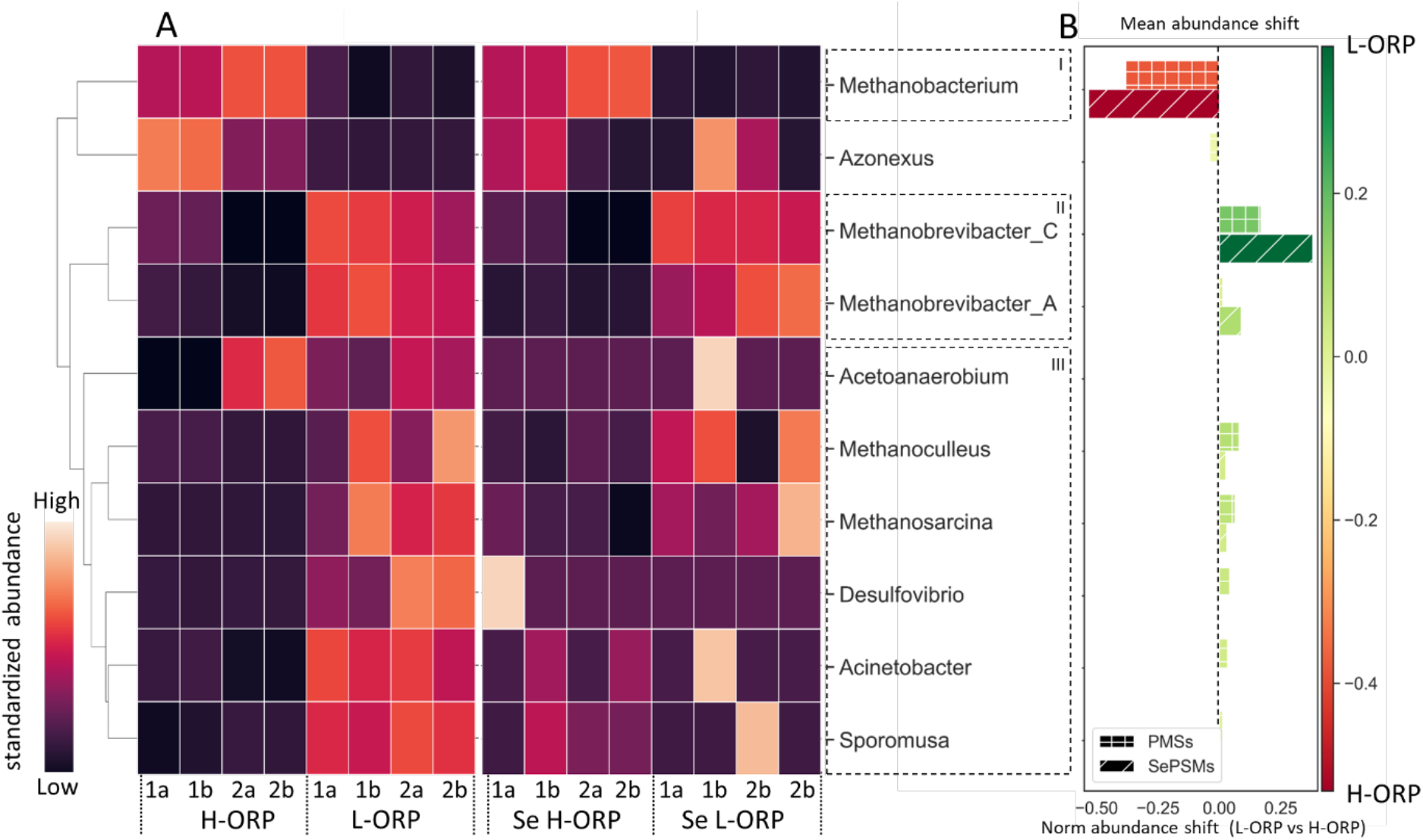
A: Clustermap of relative genus abundances in peptide spectrum matches between enrichments without reducing agent (H-ORP) and with 25mM AQDS (L-ORP) and the relative distribution of selenocysteine (Se H-ORP, Se L-ORP). Abundances were standardized across samples and clustered based on Euclidean distance. Brighter colors indicate points of higher positive abundance shift across samples for each genus B: mean shift in relative abundance of peptide spectrum matches (PSMs), as observed in (Fig 1C), and S-Se containing PSMs (SePSMs) between H-ORP and L-ORP enrichments.

The shift in taxonomic abundances between H-ORP and L-ORP enrichments displays significant shifts in relative peptide abundance and relative S-Se abundance between the two dominant methanogenic genera *Methanobacterium* and *Methanobrevibacter* C (Figure 3B). In the H-ORP environment, *Methanobacterium* saw a 39% increase in relative abundance compared to L-ORP. In contrast, the total percentage of selenocysteine peptides increased five-fold from 2.5% to 0.51%. Similarly, *Methanobrevibacter* C had 27% lower abundance in H-ORP and decreased its selenocysteine percentage from 2.24% to 1.21%. This indicates that a shift in taxonomic abundance between the two major methanogens in these samples is joined by a more substantial shift in their selenocysteine incorporation levels. Therefore, increased Se incorporation confers a competitive advantage to *Methanobacterium* and allows it to become more dominant at higher ORP (Figure 1A).

Since the importance of cysteine and selenocysteine has been established in methanogenesis, the presence of S-Se substitutions was investigated in key methanogenic enzymes of *Methanobacterium* and *Methanobrevibacter* C. This revealed strikingly high Se observations, in up to 20 % of spectra for tetrahydromethanopterin S-methyltransferase (Mtr) in both genera in the H-ORP samples (Figure 4C). Compared to the overall 2 % rate (Figure 2C) of H-ORP samples, selenocysteine is more frequently incorporated into specific methanogenic enzymes. Differing selenocysteine incorporation rates were present between *Methanobacterium* and *Methanobrevibacter* for both conditions, highlighting the dynamic nature of these putative selenocysteine incorporations (Figure 4C).

**Figure 4.**
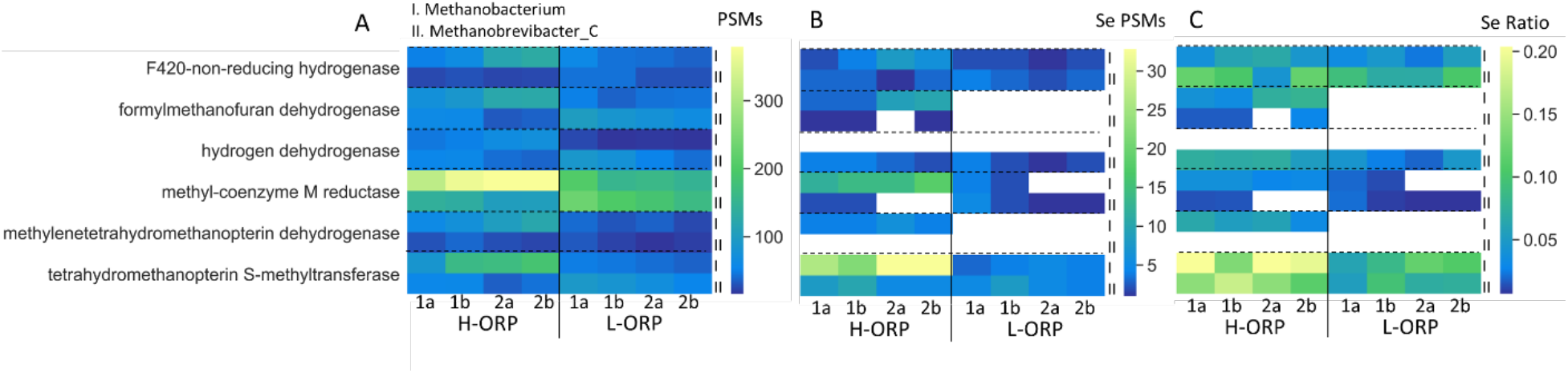
A: Peptide spectrum matches (PSMs) of key methanogenic enzymes observed for *Methanobacterium* (top row, I) and *Methanobrevibacter* C (bottom row, II), without reducing agent (H-ORP) and with 25 mM AQDS (L-ORP) B: peptide spectrum matches containing the S-Se mass shift of selenocysteine, C: the fraction of peptides that contained selenocysteine.

## Discussion

This study described the physiological response to oxidative stress by investigating the expressed proteomes under different redox conditions. Enrichment of full-scale sludge without added oxygen-scavengers in the medium yielded a tolerant community which could produce methane after being repeatedly exposed to a higher ORP (Fig. 1A). Addition of 25mM AQDS was effective at generating an optimal redox environment for methanogenesis (Fetzer 1993, Vongvichiankul 2017), and reduced AQDS acted as a redox buffer to lower ORP variability (Li 2013) (Figure 1A). AQDS has previously been studied as an additive in methanogenesis (Liang 2020, Xu 2022), and though the addition of over 20mM AQDS requires an adaptation period (Cervantes 2003, Figure S2), it should be explored as a strategy to improve oxygen tolerance.

*Methanobacterium* and *Methanobrevibacter* C dominated both communities and showed similar functional profiles. A two and a half fold increase in the ratio of selenocysteine incorporation was observed between the H-ORP and L-ORP enrichments (Figure 2C) and saw the highest incorporation in methanogenic enzymes (Figure 4). The enrichment grown in H-ORP expressed more enzymes related to mitigation of oxidative damage, as well as selenium-associated proteins (Figure 2A). By including an S-Se mass shift as modification in proteomic database search, selenocysteine incorporations could be mapped.

### Advantages of selenocysteine

Due to their higher enzymatic efficiencies than cysteine counterparts, selenocysteine enzymes have been described as super-cysteine-enzymes (Axley 1991, Rother 2010). Next to higher enzymatic activity, their oxygen tolerance (Evans 2021) and potential reversibility of oxygen-induced damage (Maroney 2018) make them suitable for micro-aerated environments. While their increased nucleophilicity and lower redox potential (Arnér 2010) could assist in the performance of methanogenesis in redox conditions that would otherwise inhibit methanogenesis (Hirano 2013). Selenocysteine incorporation could be beneficial in anaerobic digestion processes, in which operating conditions can easily cause exposure to oxygen and lead to reduced efficiency (Botheju 2011). Selenocysteine could also be of interest in systems with amendment of micro-aeration as a strategy to improve hydrolysis and methane production (Muller 2022, Shanfei 2022). An increased selenocysteine ratio coincided with increasing relative abundance in this enrichment, indicating that selenocysteine incorporation confers a competitive advantage (Figure 4) and could have enabled specific methanogens to become favoured. In this study, oxygen-sensitive Class I methanogens (*Methanobacterium* and *Methanobrevibacter*) dominated the H-ORP environments, while the L-ORP environment had increased abundance of more oxygen tolerant Class II methanogens (*Methanosarcina, Methanoculleus*). This opposes the proposed higher oxygen tolerance among Class II methanogens (Lyu 2018). Being amongst the most abundant archaea in the human gut (Gaci 2014), the oxygen tolerance of *Methanobrevibacter* and *Methanobacterium* has broader implications for the localization of methanogens within the epithelium (Tholen 2007) and adds to the importance of selenium as a nutrient for human health (Ferreira 2021).

### Selenocysteine incorporation strategies

Although selenoproteins have been observed across all domains of life, selenocysteine utilization has been described in relatively few archaea, mainly methanogens (Rother 2018, Zhang 2022), and has only recently been associated with the *Asgardarchaeota* superphylum (Liu 2018, Sun 2021). Considering the benefits of selenocysteine, it is surprising that selenium utilization is not more widespread. The presence of numerous selenocysteine incorporations combined with the lack of detected selenoproteins clashes with the stringent requirements of UGA, SECIS and specific tRNAs and instead, hint at a more flexible incorporation mechanism. SelB-like elongation factors were reported in genomes of various archaea that lacked reported selenoproteins (Atkinson 2011), and several cases of codon and tRNA flexibility exist for incorporation of selenocysteine (Schrauzer 2000, Turanov 2009, Seyhan 2015, Mukai 2016). A distinguishing feature of methanogenic archaea is their lack of direct cysteine incorporation, instead relying on SepRs and SepCysS to modify phosphoserine bound to Cys-tRNA (Sauerwald 2005, Sheppard 2008). This could imply that some archaea use similar promiscuous tRNAs that can incorporate both selenocysteine and cysteine without the need for a stop-codon and Sec-specific tRNAs (Sauerwald 2005, Bröcker 2014, Vargas-Rodriguez 2018). A similar observation was made recently for *Escherichia coli*, where it was annotated as selenocysteine “misincorporation” when selenocysteine was incorporated without UGA-codon (Hao 2020). In this case, misincorporation was deemed to be random. However, in our case, the consistently high rate of selenocysteine incorporation observed in tetrahydromethanopterin S-methyltransferase in both *Methanobacterium* and *Methanobrevibacter* C suggests otherwise and could be caused by complex regulation. Follow-up metagenomic studies would be required to verify whether selenocysteine incorporation can occur UGA-independently in these methanogenic genera and would benefit from transcriptomic analysis to track the transcription response of SECIS elements and tRNA to oxidative environments.

### Detection of Selenocysteine

While their detection seems increasingly relevant, mapping the presence of selenopeptides remains challenging. Since stop-codon recoding is not part of standard metagenomic approaches, they are underrepresented in popular databases such as GTDB. The SECIS element, which can identify selenoproteins, varies across domains and can possess both upstream and downstream locations in archaea (Zhang 2022). Additionally, due to its infrequency, selenocysteine is typically absent in homology scoring matrices, which hampers the annotation of newly identified selenopeptides. Like in phosphorylations, standard proteomic analysis often underestimates the abundance of modifications (den Ridder 2020). In the case of selenocysteine, the low pKa causes selenol groups to prefer lower pH for alkylation than is employed in standard proteomic experiments, which likely causes them to be under detected, though this also forms a target for selective selenoproteomics (Bak 2018, Guo 2018). Due to the natural isotopic abundance of selenium, selenopeptides have an altered isotopic distribution, which can form another target for their detection (Gao 2018). Lastly, *de novo* sequencing of peptides could be a promising strategy as it circumvents the issues of database construction (Kleikamp 2021), however since most sequencing tools rely on machine learning (Tran 2017), this would require retraining of models on curated spectra containing selenopeptides. Regardless, when analyzing methanogenic communities, we recommend the inclusion of S-Se mass shift as modification as a standard practice in proteomic database searches.

## Supporting information

### Initial sludge composition

Microbial community analyses of Bånlev digester sludge and transfer No. 4 of the autotrophic enrichment culture were performed via gene amplicon sequencing targeting the bacteria/archaea 16S rRNA gene variable region 4 (abV4-C) in combination with taxonomic classification against the MiDAS 4.8.1 database. DNA extraction, QC, sequencing library preparation, DNA sequencing (Illumina MiSeq 2x300 PE, scaled for 50k raw reads/sample) and data analysis with taxonomic classification were performed by DNASense ApS (Aalborg, Denmark).

**Figure SI 1:**
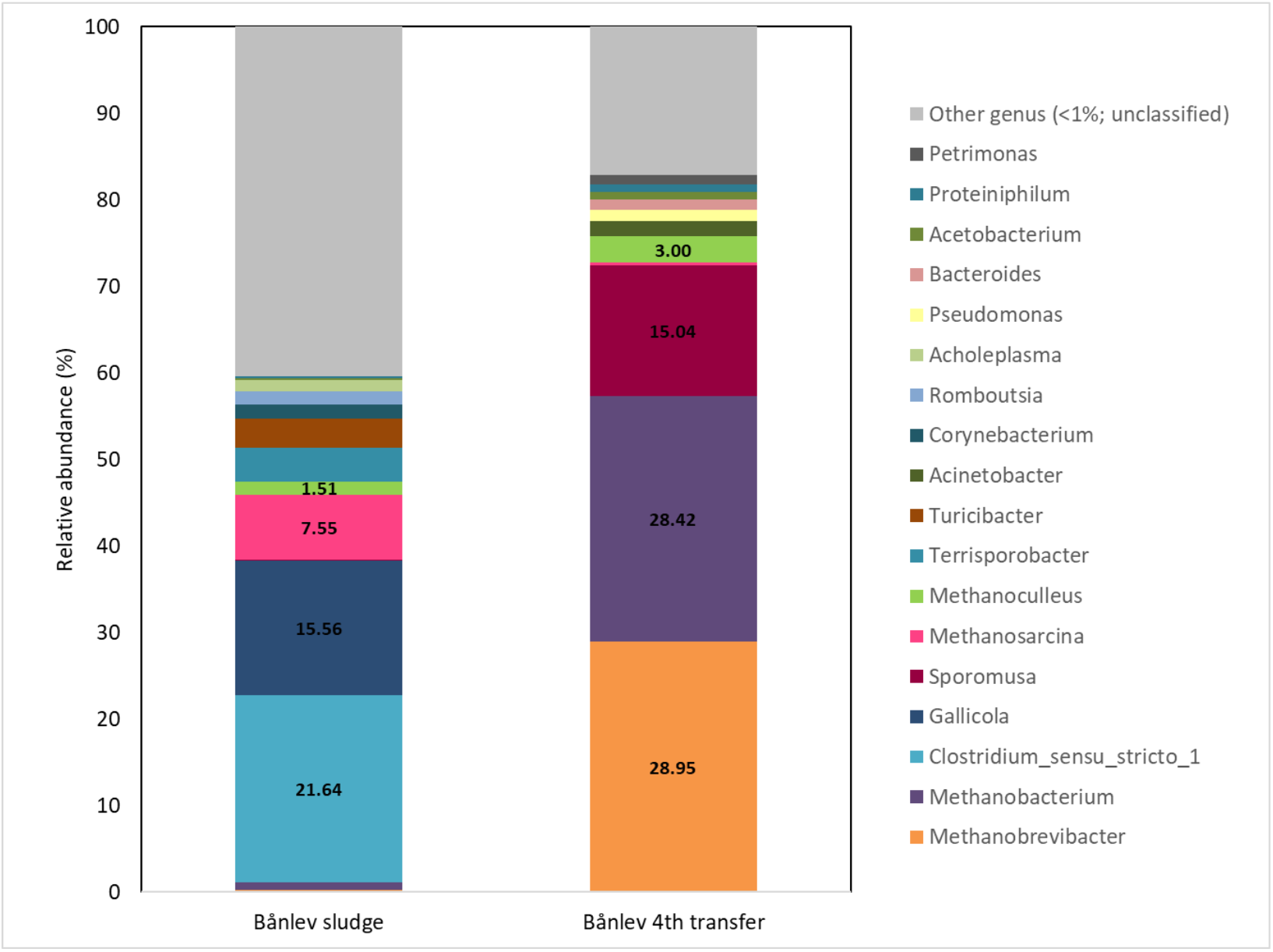
Genus level distribution from the bacteria/archaea 16S rRNA gene variable region 4 (abV4-C) from Bånlev sludge and Bånlev autotrophic enrichment (Transfer No.4).

Digestate was enriched under autotrophic conditions using 10% of inoculum and four consecutive transfers. 16s amplicon sequencing data from Bånlev sludge showed that the most abundant genera in relative abundance were *Clostridium sensu_stricto_1* (21.6%) and *Gallicola* (15.5%). *Methanosarcina* was the most abundant methanogen, with a relative abundance of 7.5%. After the four^th^ transfer, a clear difference in the microbial composition was observed since hydrogenotrophic methanogens from the genera *Methanobrevibacter* (28.95%), *Methanobacterium* (28.42%), and *Methanoculleus* (3%) were dominating the community along with homoacetogenic bacteria from the genus *Sporomusa* (15.04%).

### Analysis of methane production

Before gas sampling, pressure values from the culture’s headspace were obtained with a gas pressure sensor. The gas composition of the cultures was then monitored with a gas chromatography system (GC-2014, Shimadzu, Japan) equipped with a thermal conductivity detector (TCD) and two different columns. For CH4 analyses, a Porapak Q column (CS-Chromatographie Service GmbH, Manufacturer Item No.: 662, Langerwehe, Germany) was used with helium as carrier gas.

**Figure SI 2:**
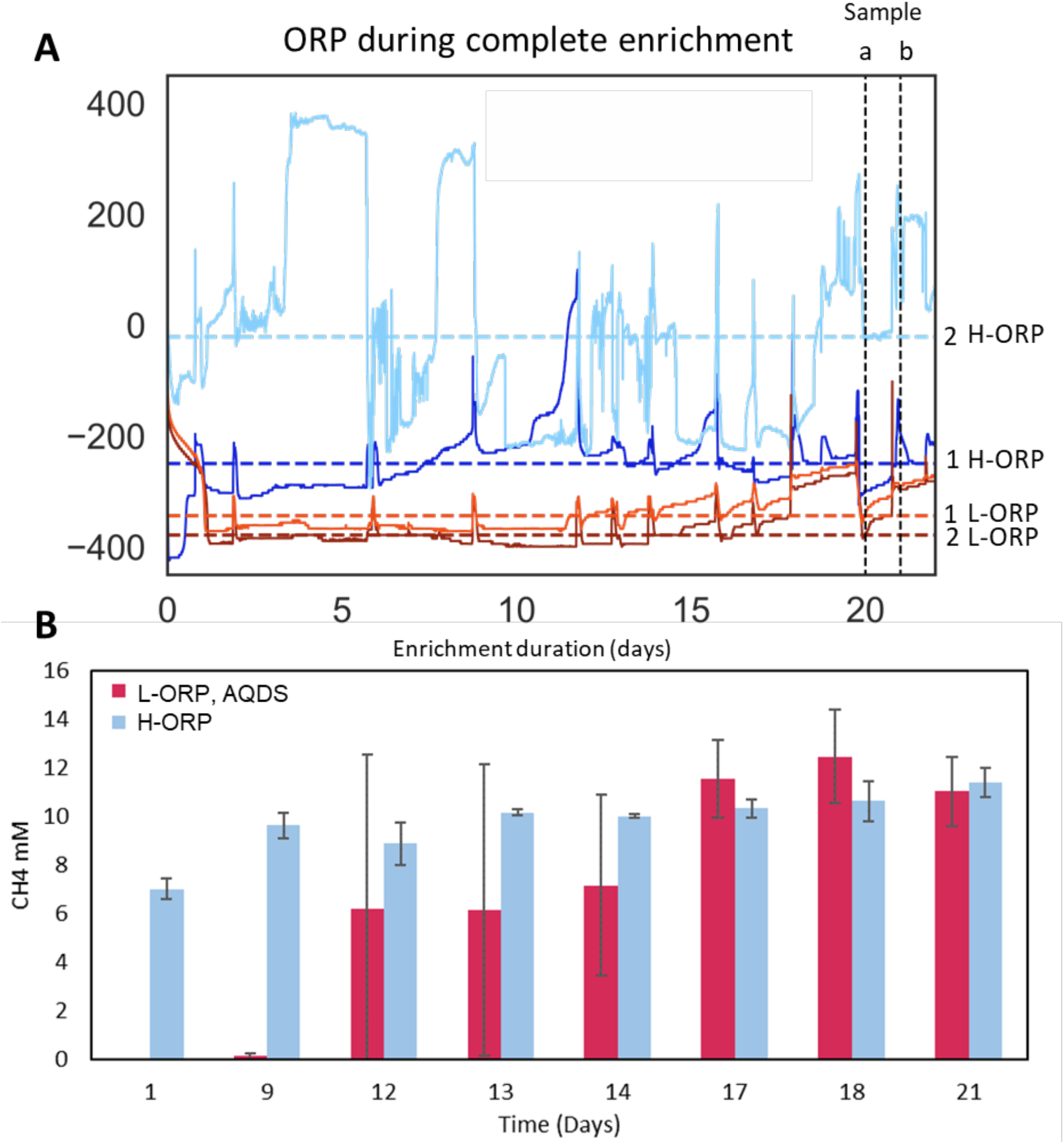
A: oxidation reduction potential (ORP) of during entire enrichment duration, of replicate (1,2) methanogenic enrichments grown with 25 mM of redox mediator AQDS (L-ORP, orange) and enrichment without reducing agents (H-ORP, blue), sampled at days 20 (a) and 21 (b) B: Evaluation of methane production during enrichment without reducing agents (H-ORP) and with 25 mM 2,7-AQDS (L-ORP)

**Figure SI 3:**
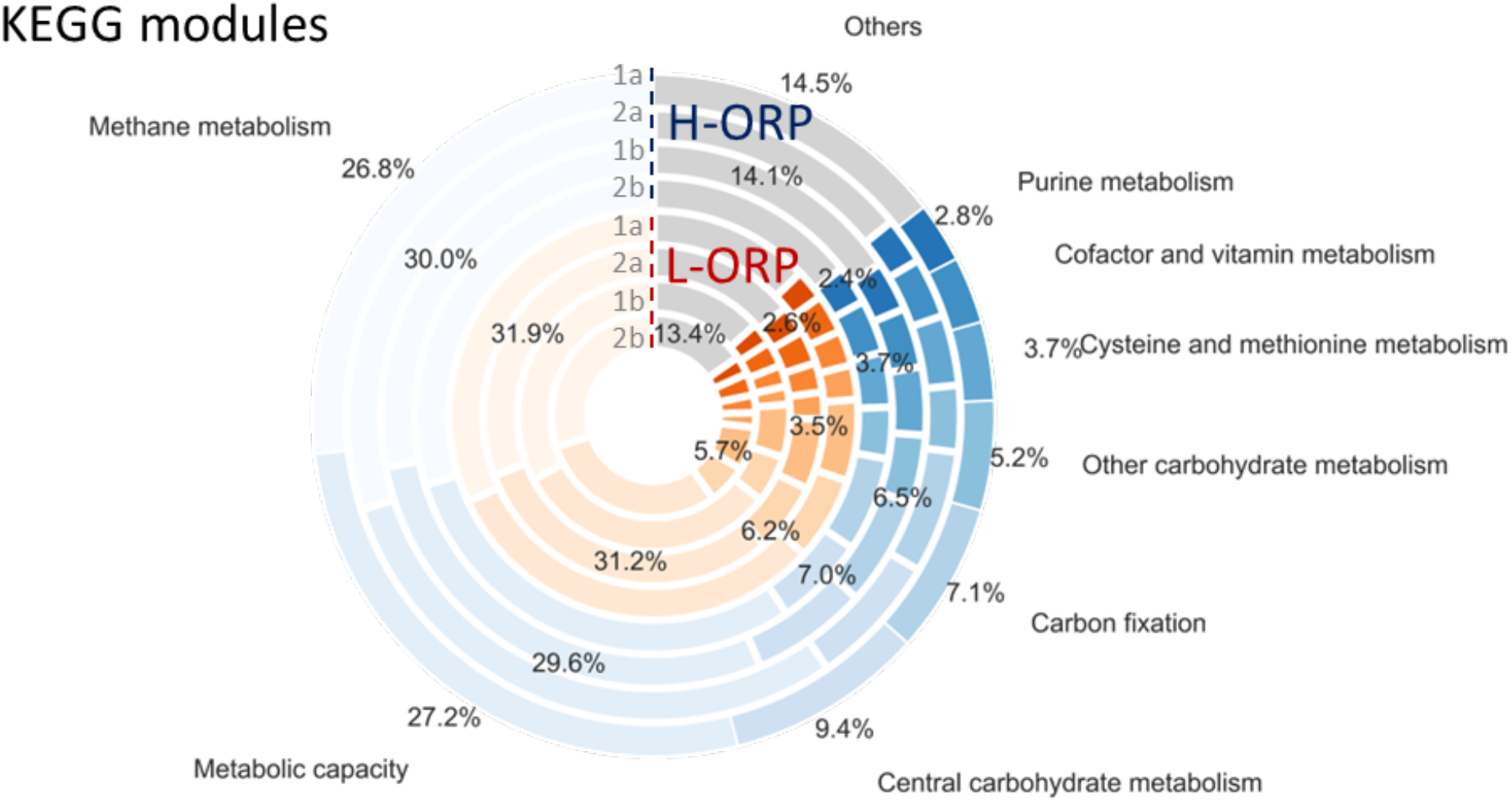
Ten most abundant KEGG modules from metaproteomic analysis, displaying the metabolism of two enrichments (1,2) without reducing agents (H-ORP) and with 25 mM 2,7-AQDS (L-ORP), sampled at days 20 (a) and 21 (b)

**Figure SI 4.**
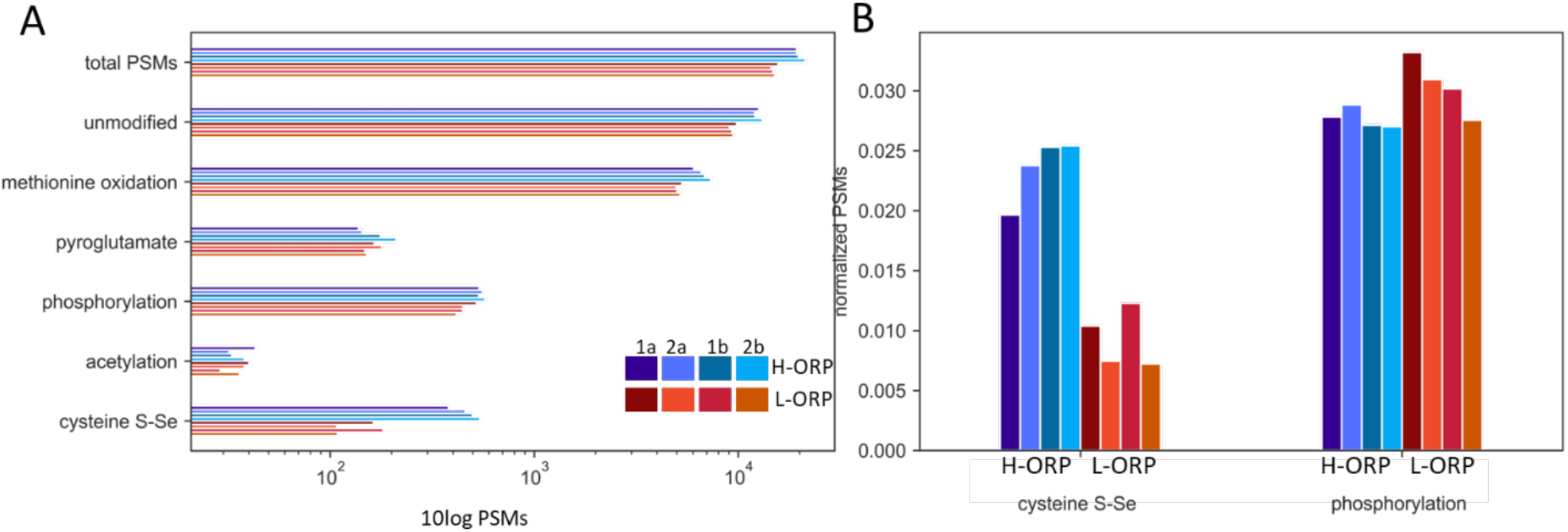
Comprehensive comparison of observed S-Se mass shift compared to other variable modifications in enrichments without reducing agents (H-ORP) and with 25 mM 2,7-AQDS (L-ORP), with technical replicates (a,b) and biological replicates (1,2). A: absolute number of peptide spectrum matches (PSMs) associated with common modifications, B: close-up of normalized PSMs of selenocysteine incorporation compared to phosphorylations.

**Figure SI 5:**
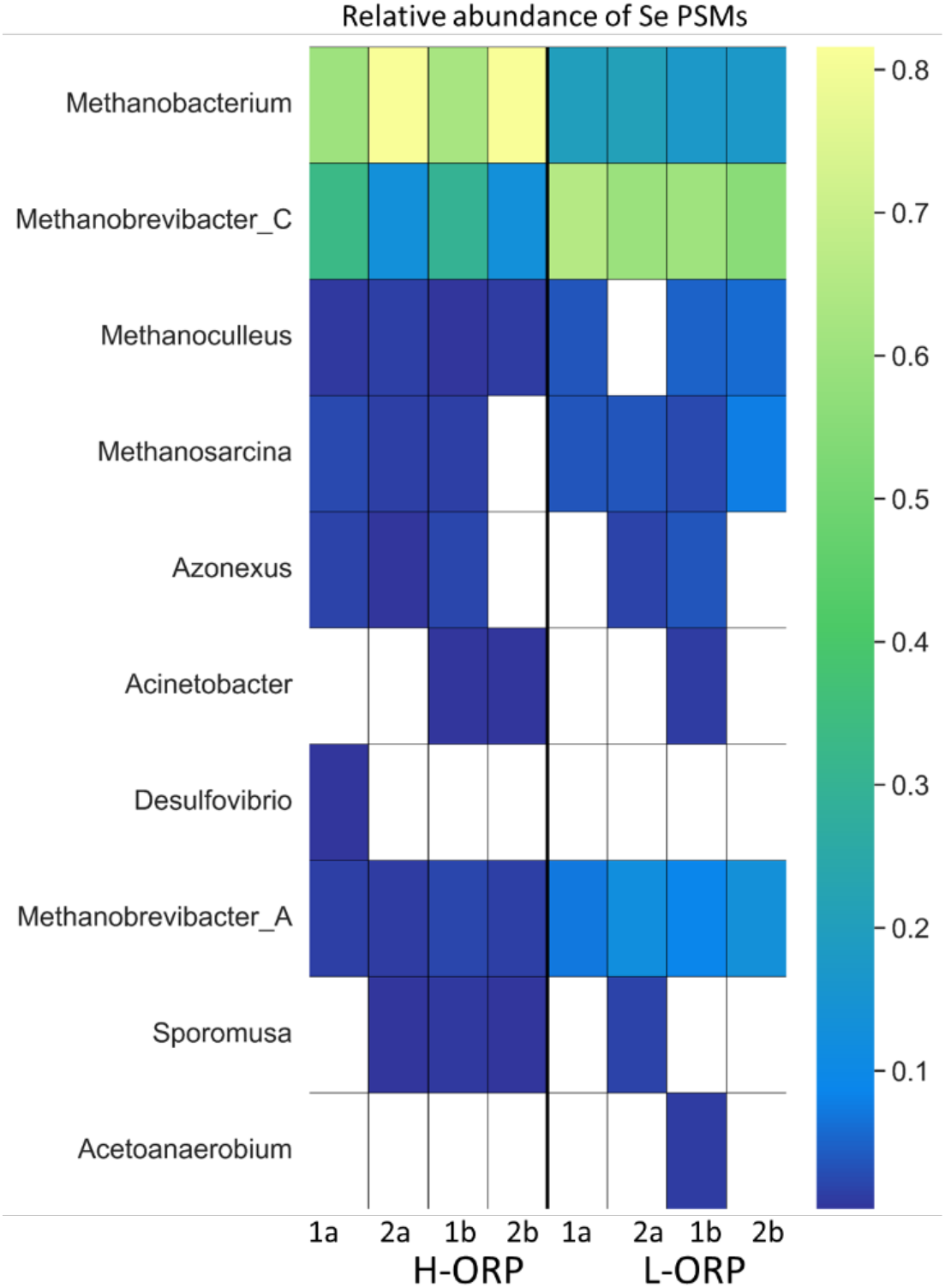
relative abundance of observed selenocysteine peptide spectrum matches (PSMs) for the ten most abundant genera in enrichments without reducing agents (H-ORP) and with 25 mM 2,7-AQDS (L-ORP), with technical replicates (a,b) and biological replicates (1,2)

**Figure SI 6:**
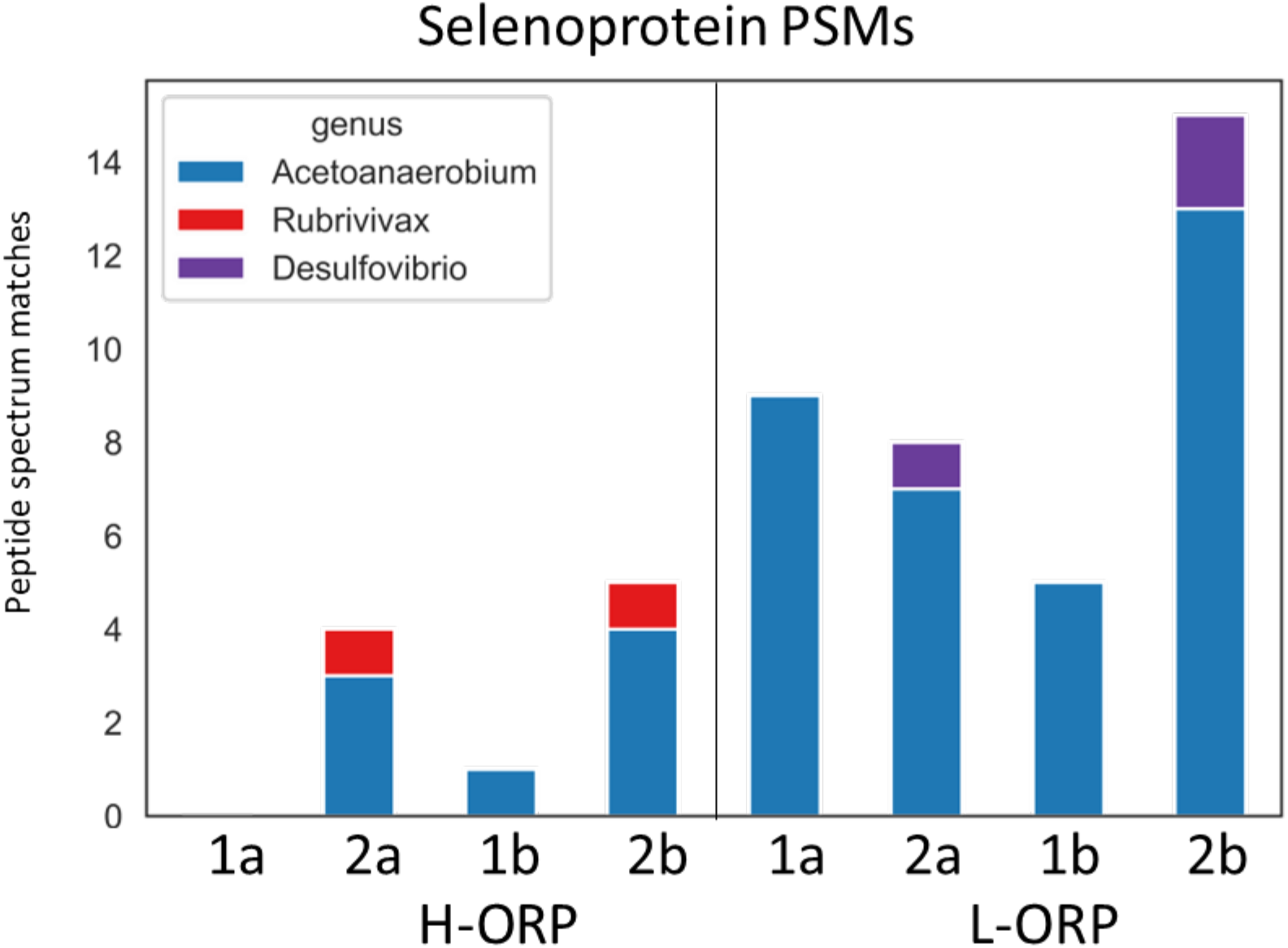
PSMs of peptides belonging to known selenoproteins, as annotated with UniprotKB reference proteomes in enrichments without reducing agents (H-ORP) and with 25 mM 2,7-AQDS (L-ORP), with technical replicates (a,b) and biological replicates (1,2)

**Table SI 1:** Functional targets used to Identify proteins involved in ROS mitigation.

| Name | Function | Database | Accessions |
| --- | --- | --- | --- |
| disulfide reductase | damage repair | EC | 1.8.4.1 1.8.4.10 1.8.4.11 1.8.4.12 1.8.4.13 1.8.4.14 1.8.4.15 1.8.4.16 1.8.4.2 1.8.4.3 1.8.4.4 1.8.4.5 1.8.4.6 1.8.4.7 1.8.4.8 1.8.4.9 1.8.4.B3 |
| superoxide reductase<br>dismutase | O2 elimination | EC | 1.15.1.1 1.15.1.2 |
| F420 H2 oxidase | O2 elimination | EC | 1.5.3.22 |
| catalase | O2 elimination | EC | 1.11.1.6 1.11.1.21 |
| peroxidase | O2 elimination | EC | 1.11.1.1 1.11.1.B2 1.11.1.2 1.11.1.3 1.11.1.5 1.11.1.B6 1.11.1.B7 1.11.1.7 1.11.1.B8 1.11.1.8 1.11.1.9 1.11.1.10 1.11.1.B10 1.11.1.11 1.11.1.12 1.11.1.13 1.11.1.14 1.11.1.16 1.11.1.17 1.11.1.18 1.11.1.19 1.11.1.21 1.11.2.2 1.12.1.1 |
| glutathione metabolism | O2 elimination | KEGG | K11204 K11205 K01919 K21456 K01920 |
| cysteine and methionine<br>metabolism | O2 elimination | KEGG | K11358 K13034 K05810 K01251 K23370 K20249 K00548 K00831 K23975 K13061 K21623 K00549 K10150 K01505 K01758 K14455 K00827 K09758 K01762 K17462 K00024 K20250 K21456 K00640 K00812 K00641 K22847 K00544 K11204 K01738 K01740 K01752 K01566 K00797 K01919 K08969 K23976 K03334 K01767 K05396 K16843 K18284 K00003 K22954 K24034 K16054 K25316 K19696 K01011 K00813 K17399 K22956 K00558 K00838 K22846 K08963 K25035 K16846 K17216 K10764 |
|  |  |  | K00025 K00547 K13060 K12526 K00651 K00837 K12524 K24042 K01697 K13062<br>K00826 K17069 K00899 K23977 K09880 K01739 K22968 K20772 K00058 K01761<br>K20021 K25317 K08964 K12339 K22955 K01920 K00789 K00816 K14155 K22957<br>K00026 K22207 K00016 K12525 K01244 K00811 K00772 K00928 K00832 K17950<br>K23304 K00802 K07173 K08967 K00456 K16845 K12960 K01611 K08966 K05953<br>K00815 K00133 K20248 K01243 K05933 K16844 K17217 K11205 K01760 K14454<br>K08968 K17989 K08965 K17398 |
| selenocompound metabolism | Selenium | KEGG | K01763 K00957 K00384 K01042 K17050 K17052 K12527 K22182 K24042 K00548<br>K00549 K01874 K00958 K00955 K17051 K01758 K01739 K07309 K13811 K01761<br>K08247 K12528 K00562 K12529 K01760 K10837 K00816 K14155 K11717 K03341<br>K00956 K01008 K07310 |
| Base excision repair | damage repair | KEGG | K07462 K03506 K10563 K01151 K03505 K01972 K10803 K10801 K03512 K02324<br>K10771 K10776 K24070 K02335 K10802 K05522 K01247 K01246 K02326 K20813<br>K03649 K10800 K03575 K13529 K03660 K10568 K04799 K10773 K02327 K03504<br>K02330 K10747 K10798 K02325 K01142 K03648 K02328 K10567 K03652 K10569<br>K21929 K04802 K10772 |
| Proteins involved in Fe-S cluster biogenesis | damage repair | KEGG | K04487 K22068 K22069 K19054 K22070 K22071 K18914 K04043 K04082 K14001<br>K07390 K22063 K22072 K22073 K22066 K22075 K23500 K23501 K23502 K23503 |
| Ferritin-like superfamily | O2 elimination | IPR | IPR009078 |
| Thioredoxin-like superfamily | O2 elimination | IPR | IPR036249 |
| Rubredoxin-like domain | O2 elimination | IPR | IPR024934 |
| Selenium associated | Selenium | IPR | IPR025862 IPR017695 IPR021603 IPR017704 IPR026366 IPR008826 IPR019870<br>IPR017697 IPR017587 IPR019872 IPR018319 IPR004534 IPR038549 IPR017584<br>IPR007672 IPR007671 IPR017700 IPR022474 IPR017701 IPR041085 IPR017698<br>IPR030805 IPR037941 IPR023061 IPR004536 IPR006338 IPR019389 IPR040434<br>IPR017582 IPR004535 IPR010970 IPR008829 IPR011893 |

